# Using computational theory to constrain statistical models of neural data

**DOI:** 10.1101/104737

**Authors:** Scott W. Linderman, Samuel J. Gershman

## Abstract

Computational neuroscience is, to first order, dominated by two approaches: the “bottom-up” approach, which searches for statistical patterns in large-scale neural recordings, and the “top-down” approach, which begins with a theory of computation and considers plausible neural implementations. While this division is not clear-cut, we argue that these approaches should be much more intimately linked. From a Bayesian perspective, computational theories provide constrained prior distributions on neural data—albeit highly sophisticated ones. By connecting theory to observation via a probabilistic model, we provide the link necessary to test, evaluate, and revise our theories in a data-driven and statistically rigorous fashion. This review highlights examples of this theory-driven pipeline for neural data analysis in recent literature and illustrates it with a worked example based on the temporal difference learning model of dopamine.

## Introduction

The statistical toolbox for neuroscience has been steadily growing in sophistication—relaxing restrictive assumptions, increasing expressiveness, and enhancing computational efficiency. These advances have enabled a recent blossoming of “data-driven” approaches to neuroscience, which aim to provide insight into neural mechanisms without testing specific computational theories. Data-driven approaches are appealing, at least in principle, for several reasons: they do not require the scientist to explicitly specify a set of hypotheses, they are unprejudiced by the scientist’s theoretical dispositions, and they avoid the problem that many computational theories are too abstract to make direct contact with neural data.

In this paper, we argue that such faith in data-driven approaches is misplaced. Far from escaping the explicit specification of hypotheses, any statistical model of neural data inevitably makes assumptions about the structure of the data, and there is no principled distinction between statistical assumptions and scientific hypotheses. (Admittedly, a purely data-driven approach is something of a straw-man, but we pursue this line of argument for pedagogical purposes). A corollary of this point is that theoretical dispositions are inescapable: it is impossible to specify a statistical model without making assumptions. The question then becomes what assumptions to make. We argue that these assumptions should be derived from computational theories, coupled with flexible statistical parametrizations that compensate for inaccuracy and under-specification of the theories.

We illustrate this argument with a worked example, using a paradigmatic neurocomputational theory: the temporal difference learning model of dopamine. We show how the computational theory can be augmented with modern statistical tools to produce a powerful data analysis methodology. This approach generates a more complete and flexible specification of the theory. Moreover, we show that this approach offers insights into the mechanisms underlying neural data that are inaccessible to purely data-driven approaches.

**Figure 1:**
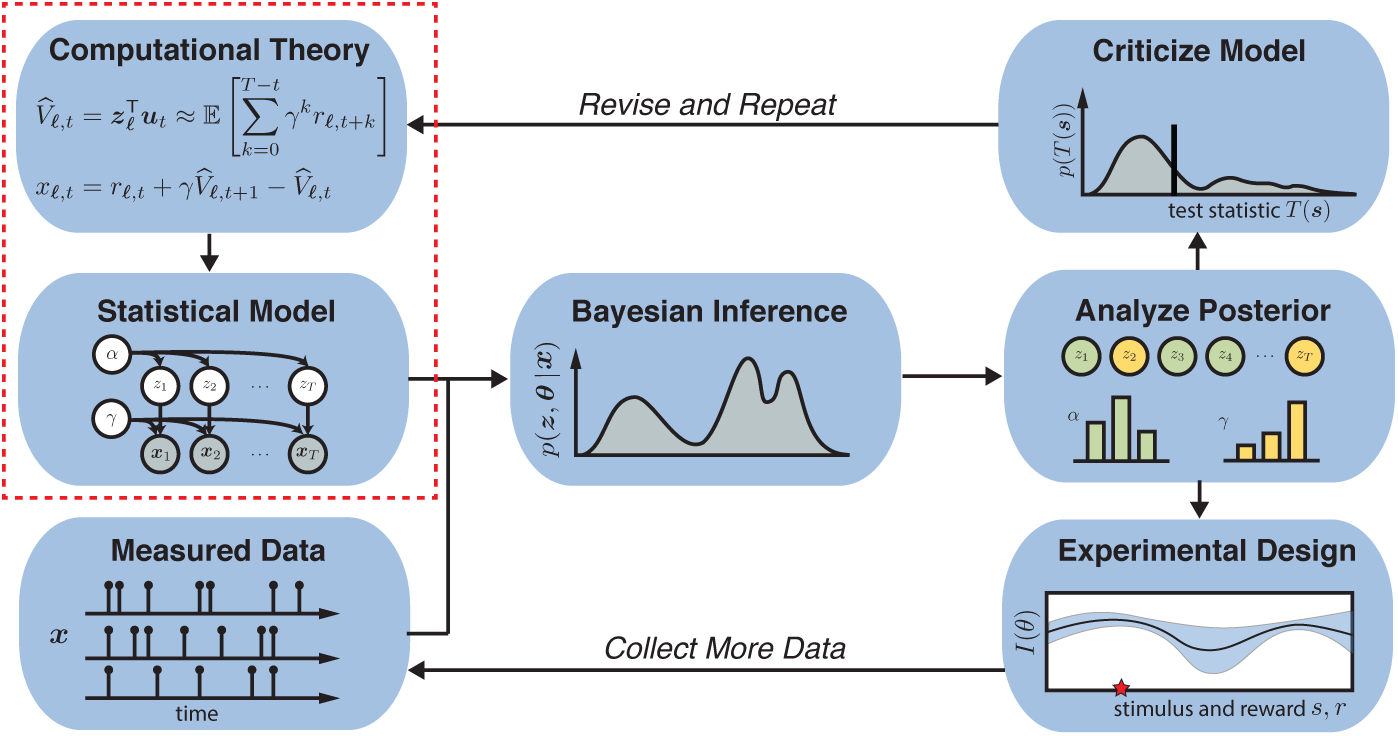
A theory-driven pipeline for neural data analysis based on “Box’s Loop”[1, 2]. This review illustrates many examples of translating theory into statistical model (red box). The benefits are many. Given a model, we may leverage a powerful toolbox of statistical techniques for inference, model criticism, and experimental design. Equally important, theory constrains the space of models and provides a critical lens through which to interpret the posterior. We will discuss advances in each stage of this pipeline.

## A Theory-driven Pipeline for Neural Data Analysis

Neural data analysis is an iterative process that begins with a data set and an idea of the underlying processes that shaped it. The first step, and arguably the most important one, is to turn that idea into a model. With a model in hand, we fit it to the data and investigate the learned parameters, searching for patterns that shed new light on the system under study. But the process does not end here; we then interrogate our model, see where it captures the data well and where it fails, and use these criticisms to suggest model enhancements or subsequent experiments. Thus, model criticism leads to a new model and another iteration of the process.

Statisticians have formalized and automated many pieces of this pipeline: models are joint distributions over data, latent variables and parameters; “fitting” is performed by posterior inference; criticism is carried out with statistical tests; and optimal experimental design suggests what experiment to run next. This cyclic process of probabilistic modeling, inference, and statistical criticism is known as “Box’s loop” [1, 2], and later sections of this review will discuss many recent advances in each stage of the pipeline.

Still, the art of carving a manageable class of models from the infinite space of possibilities remains the province of the practitioner. It is here that computational theory can play a vital role, since theories suggest what structure and patterns may exist in the data. In doing so, theories constrain the class of models and make it easier to search, and provide a lens through which to interpret model parameters. These benefits are reciprocated: once a theory has been translated into a probabilistic model, a vast statistical toolbox can be harnessed to test and refine it in light of data.

Theory-driven statistical models are the norm in many fields, most notably in physics, where strong quantitative predictions can be derived from first principles. For example, the discovery of the Higgs boson relied on statistical tests based on predictions of the standard model [3]. Perhaps it is unsurprising, then, that some of the best examples of theory-driven statistical analyses in neuroscience arise from detailed, biophysical models of single cells. For example, Huys and Paninski [4] use the Hodgkin-Huxley model to derive a probabilistic model for noisy membrane potential recordings. The conductances of various ion channels are free parameters of their model, and the time-varying channel activations are their latent states. Given the membrane potential, their goal is to infer the conductances, integrating over possible activation states. The highly nonlinear nature of the Hodgkin-Huxley dynamics and the potentially large number of different channel types present a formidable challenge, but biophysical constraints limit the space of feasible parameters. In recent work, these methods have been extended to data in which only spike trains are observed [5], which present an even greater challenge.

Many models in neuroscience are phenomenological rather than mechanistic in nature. One step up from biophysical models are firing rate models like the generalized linear model (GLM) [6, 7, 8]. Recent work has extended these classical models to make them more flexible [9], more biophysically inspired [10], and more interpretable [11]. While the GLM omits many mechanistic details, in fully-observed networks its weights can be roughly interpreted as synaptic strengths [12, 13]. However, the weights of the standard GLM are static, even though synaptic plasticity may be at work in many neural recordings. While the space of all possible dynamic GLM’s is intractably large, theories of synaptic plasticity place strong constraints on how synaptic weights evolve over time in response to preceding activity. A number of authors have leveraged these constraints to develop theory-driven GLM’s with time-varying weights and have shown how alternative models of synaptic plasticity can be compared on the basis of their fit to spike train data [14, 15, 16].

This approach extends to computational theories as well, and is exemplified in the work of Latimer et al. [17]. The authors reconsider the long-standing theory of evidence accumulation in lateral intraparietal (LIP) cortex [18], and ask whether patterns that emerge in trial-averaged data are borne out in individual trials. Specifically, do the firing rates of neurons in LIP slowly ramp as evidence is accumulated, or do they exhibit a discrete jump in firing rate? Theory suggests the former, whereas the latter would indicate that LIP may not be the site of integration. Critically, both theories would yield the appearance of a ramp in trial-averaged firing rate. Latimer et al. [17] formulate both theories as probabilistic models for single trial data, fit these models with Bayesian inference, compare them on the basis of the marginal likelihood of the data, and find that a large fraction of neurons are better explained by the discrete jump model. This provides statistical evidence with which to assess and reevaluate canonical theory. Indeed, this work has prompted further assessments of their modeling assumptions and the validity of their conclusions [19]—a prime example of Box’s loop in action post-publication.

Integrative approaches to computational theory and statistical analysis have also been pursued in higher-level cognition. Detre and colleagues [20] used Bayesian inference to identify a nonmonotonic relationship between memory activation (as measured by functional MRI) and subsequent memory, as predicted by a competition-dependent theory of episodic memory [21]. The same analytical approach was used to identify other nonmonotonic effects of retrieval strength on memory [22, 23].

The aforementioned examples stand in contrast to many dimensionality reduction methods like PCA, tSNE [24], and others [25], and differ as well from general-purpose state space models [26, 27, 28] and recurrent neural network models [e.g. 29] for neural data. Such methods start with very weak assumptions—linear embeddings or low-dimensional dynamics—and, in this sense, allow the data to speak freely. Thus, they are invaluable exploratory tools. However, in the absence of a theory, the inferred low-dimensional states and projections require careful interpretation. In many cases, theories correspond to special cases of these general-purpose models, and thus help address issues of interpretability.

The landscape of neural data-analysis is not as strictly divided into top-down and bottom-up approaches as the preceding discussion may suggest. Indeed, many models fall somewhere in the middle, incorporating aspects of theory while allowing flexibility in aspects that are less certain. Wiltschko et al. [30] strike such a balance in their model for depth videos of freely behaving mice. Starting with the classic ethological theory that behavior is composed of a sequence of discrete, resuable units, or “syllables,” the authors propose an autoregressive hidden Markov model to discover these syllables from raw data. However, since the number of syllables is not known *a priori*, the authors use a Bayesian nonparametric prior distribution [31] to determine the number of states in a data-driven manner.

These works exhibit a diverse array of “theory-driven” neural data analyses, but the best way to understand this pipeline is through an example.

## A Worked Example

There is no single recipe for translating computational theories into probabilistic models of data, but the conversion necessarily involves answering a few basic questions. Which theoretical variables and parameters are observed and which are latent? How are they encoded by the neural system under study? How do these variables evolve over time? What are the sources of noise in the system and in the measurements? The answers to these questions inform statistical models of data that in turn define distributions of likely patterns of neural activity. We will illustrate this translation with a simple worked example^1^.

Temporal difference (TD) learning [33] is a classical algorithm by which agents, over the course of many trials, learn to use sensory cues to predict the discounted sum of future rewards. Assume that there are *L* trials, each lasting *T* time steps. On trial ℓ the agent receives a sequence of stimuli, which are stored and encoded as vectors, *u_ℓ,t_* and a corresponding sequence of rewards, *r_ℓ,1_*…,*r_ℓ,T_*, most of which may be zero. In a classical conditioning experiment, the stimulus may be a light at time *t* followed by a reward some number of time steps in the future, and *u_ℓ,t_* may encode, for example, the number of time steps since the bell was heard. The agent then uses this encoding to compute a *value function* for the given trial and time step,

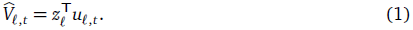

In reinforcement learning, the value is the total amount of future reward to be expected after receiving input *u_ℓ,t_* However, according to the theory, the reward is discounted by how long one must wait before receiving it. For example, a reward *k* time steps into the future is down-weighted by a factor of γ^*k*^, where γ∈ [0, 1] is the *discount factor*. The agent’s goal is to adjust the weights^2^ of its value function z_ℓ_, such that the value function approximates this discounted sum of expected future rewards,
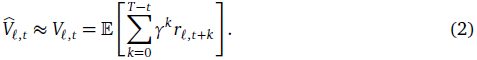

If the environment is a Markov decision process, the target value function can be written recursively as 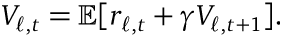 When the value function equals the cumulative discounted reward, the *reward prediction error*,

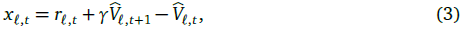
will equal zero. Intuitively, the reward prediction error provides an instantaneous estimate of how well the value function predicts the received reward. Thus, to improve its value function, the agent should adjust its weights to reduce this error. Indeed, this is accomplished by the simple learning rule, 
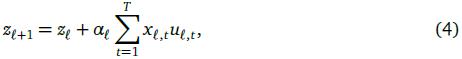
which can be seen as a form of stochastic gradient descent on the (squared) reward prediction error with learning rates α_1_,…, α_*L*_. In the following experiments, we will consider two learning schedules: a power-law schedule,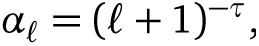and a constant schedule, α_ℓ_≡τ. In both cases, assume τ ∈ [0, 1].

Schultz et al. [34] found that the firing rates of dopaminergic neurons in the ventral tegmental area (VTA) mimic the reward prediction errors essential to the TD-learning algorithm. Moreover, it is hypothesized that cortex represents the stimulus, striatum represents the value function estimate, and VTA activity modulates plasticity of synapses from cortex to striatum [35]. Still, many important questions remain, like how learning schedules, which affect this plasticity, vary from trial to trial in real neural circuits. As a didactic exercise, we will we use the TD learning theory to construct a probabilistic model for neural data, and use that model to compare between different learning schedules in a statistically rigorous manner.

Suppose that we have access to simultaneous noisy recordings of a VTA neuron and an upstream population of *N* cortical neurons. As has been hypothesized, we will assume the VTA neuron encodes reward prediction error, χ_ℓ,t_ and the cortical neurons carry the stimulus encoding, *u_ℓ,t_*. Moreover, assume we know the reward signal, *r*_ℓ,t_.According to the TD learning theory, the cortical and VTA signals are related via a value function, which is determined by an unobserved and dynamic set of weights at each trial. In other words, the theory implies that the reward prediction errors follow a latent state space model whose hidden states are the weights, *z*_ℓ_, and whose parameters vary from trial to trial according to the cortical inputs, rewards, and prediction errors. If we assume Gaussian noise in the weight updates and observations, the theory implies that the VTA activity follows a Gaussian linear dynamical system (LDS) with non-stationary parameters.

To see this equivalence, we rewrite the TD learning updates in standard state space notation:

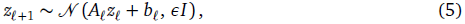

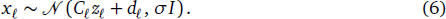

Here, the latent states are the weights, 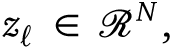 and their dynamics are determined by, *A_ℓ_=I* and 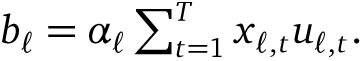 That is, the weights follow a random walk biased by the learning rate, error signal, and inputs. The emissions are vectors of observed VTA activity, *x*_*ℓ*_ = [*x*_*ℓ*_,1; …; *x*_*ℓ*_,_T−1_], and they are determined by the matrix *C*_*ℓ*_ = [*c*^T^_*ℓ*, 1_; …; *c*^T^ _*ℓ*,*T*−1_], where *c*_*ℓ,t*_ = *γu*_*ℓ*_,_*t*+1 −_ *u*_*ℓ,t*_, and by the bias vector, *d*_*ℓ*_ = [*d*_*ℓ*,1_, …, *d*_*ℓ*_,_T−1_], where, *d*_ℓ,*t*_=*r_ℓ,t+1_* Note that both the dynamics and emission parameters are non-stationary; that is, they vary from trial to trial. The noise in the weight updates is governed by ε, and the noise in the observations is governed by σ. Referring back to equations (1)–(4), we see that the exact TD learning model is recovered in the noise-free limit. The free parameters are θ=(τ, γ, ∈,σ,)—the learning rate parameters, discount factor, and noise variances.

We call this constrained model a temporal difference LDS (TD-LDS). Importantly, by translating the TD learning theory into a constrained Gaussian LDS, we have reduced it to an essentially solved model with very mature estimation and interpretation procedures [36]. In the next section we will show how to infer the states and parameters of the TD-LDS from data.

What assumptions did we make in deriving the TD-LDS? First, we assumed Gaussian noise in both the observed reward prediction errors and the weight dynamics. If we observed spike counts instead, the resulting model would be more akin to a Poisson linear dynamical system (PLDS) 26, 27]. If we had assumed a nonlinear model for the value function, i.e V_*ℓ,t*_ = *f*(*z*_*ℓ*_ *u*_*ℓt*_), then both the dynamics and observation models would be nonlinear in *z*_ℓ_, which would necessitate more sophisticated inference procedures. We will only consider the linear Gaussian case in this didactic example.

## Bayesian Inference

Bayesian inference algorithms take as input the observed data, *x*, and a probabilistic model, *p*(*x*, *z*, θ), and output the posterior distribution over the latent variables and param-eters of the model, *p*(*z*,θ | *x*). By Bayes’ rule, this posterior distribution is given by,

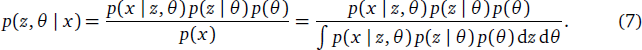

With this posterior distribution in hand, we can answer a host of scientific questions. We can estimate the posterior mean and mode (the maximum *a posteriori* estimate), and we can provide Bayesian credible intervals by computing the quantiles of the posterior distribution. Moreover, we can predict what future data would look like with the *posterior predictive* distribution,

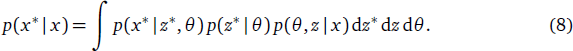

which integrates over the space of parameters and latent variables, weighting them by their posterior probability given the data seen thus far. As we will show below, these functions of the posterior distribution provide principled means of comparing and checking models.

Unfortunately, the normalizing constant on the right-hand side of Bayes’ rule, *p*(*x*), also known as the *marginal likelihood*, requires an integral over all possible parameters. This integral is intractable for all but the simplest models, so in practice we must resort to approximate techniques like Markov chain Monte Carlo (MCMC) [37] or variational inference [38, 39]. MCMC algorithms approximate the posterior distribution with a collection of samples collected by a Markov chain that randomly walks over the space of parameters. With a carefully tuned random walk, the stationary distribution of the Markov chain is equal to the desired posterior distribution so that, once the chain has converged, parameters are visited according to their posterior probability. In contrast, variational inference algorithms specify a family of “simpler” distributions and search for the member of this family that best approximates the desired posterior. Thus, they convert an integration problem of computing the denominator of Bayes’ rule into an optimization problem of searching over the variational family. Of course, both approaches present challenges—how to tell if a Markov chain has converged? how to select and search over a variational family and diagnose errors in the obtained approximation?—making Bayesian inference both an art and a science.

Fortunately for the practitioner, as probabilistic programming packages grow in sophistication, the nuances of approximate inference play a lesser role. Probabilistic programming languages like Anglican [40], Stan [41], Venture [42], and Edward [43] remove the burden of deriving and implementing an inference algorithm, and simply require the practitioner to specify their probabilistic model and supply their data. Under the hood, these packages automatically derive suitable MCMC or variational inference algorithms. In practice, some care must be taken to ensure these systems provide accurate inferences, and these tools still cannot compete with well-tuned, model-specific inference algorithms. However, they can dramatically accelerate the scientific process by enabling rapid iteration over models. Once a model has been selected, time may be invested in deriving bespoke inference algorithms for peak performance.

We have taken an intermediate approach to inference in our working example. After reducing TD learning theory to a canonical state space model, we leverage off-the-shelf inference algorithms for the latent states and develop model-specific updates only for the parameters. Specifically, given the discount factor and the learning schedule, the posterior distribution over latent states is found with a standard message passing algorithm [39]. Given a distribution over latent states, we estimate the most likely learning schedule parameters and discount factor with hand-derived updates. We alternate these two steps— updating the latent states and re-estimating the parameters—in our variational inference algorithm.

Figure 2 illustrates some of the results of our Bayesian inference algorithm. Panel (e) shows the posterior mean of the states, which in this model correspond to the weights of the value function. From the posterior distribution over weights, we derive the distribution over the value function, which is linear in the weights (c.f. (1)). Panel (f) shows the true and inferred value function at early (blue), middle (red), and late (yellow) trials, along with the uncertainty under the posterior. Likewise, panel (g) shows the inferred learning rate under two different models: a model with constant rates and a model with rates that decay according to a power law (the true model in this case). Posterior visualizations like these play a critical role in the scientific process, providing views of the low-dimensional structure of complex data. However, these visualizations are only useful to the extent that the model captures meaningful structure. Panel (h) exemplifies this point: a standard LDS with the same latent dimension as the TD-LDS provides a very good fit to the data, but its latent states look like pure noise. Without a theoretical structure with which to interpret this low-dimensional projection, the latent states are meaningless.

**Figure 2:**
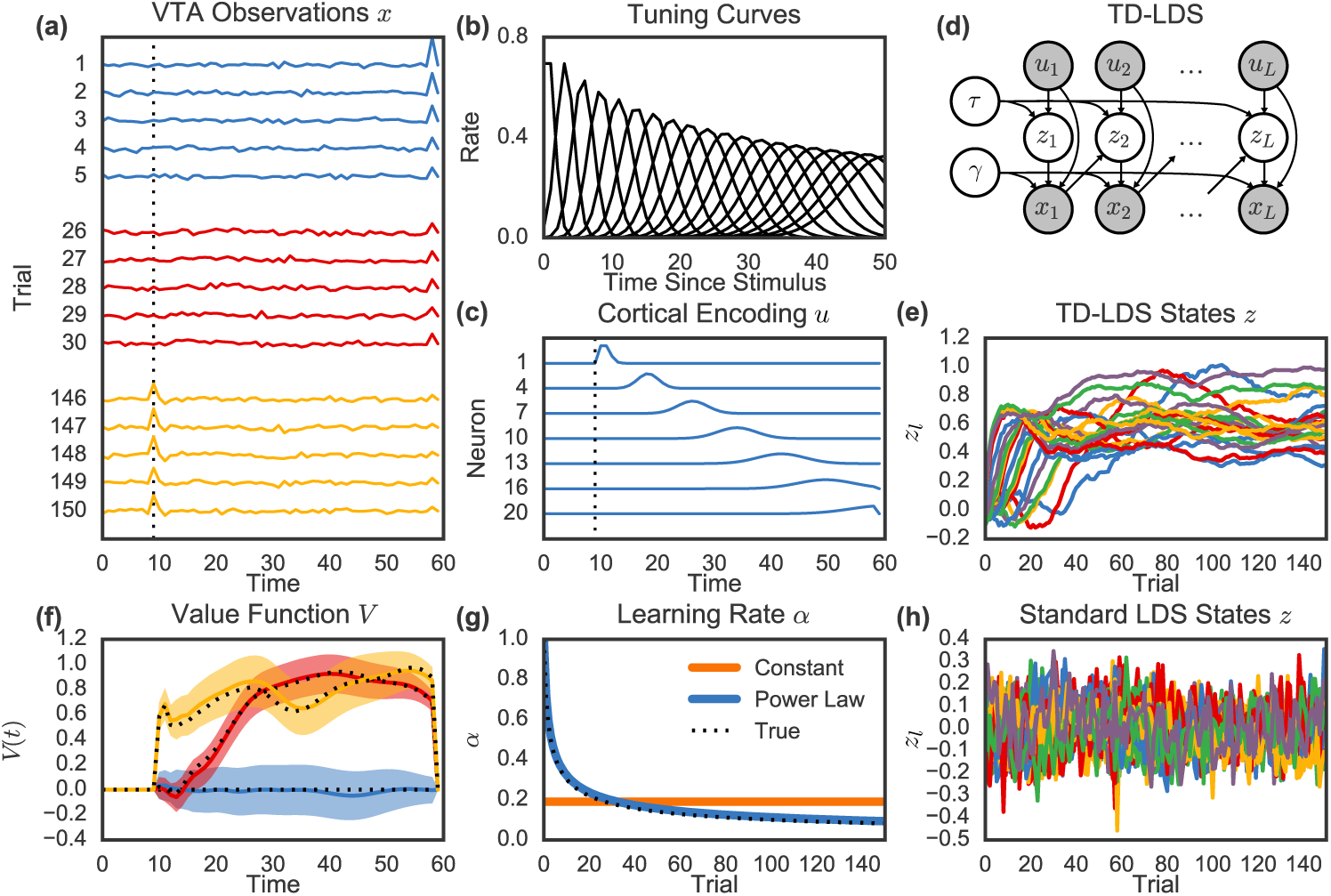
An illustrative example of using the theory of TD learning to constrain a probabilistic state space model for neural data. **(a)** Simulated example of a dopamine neuron encoding reward prediction error in VTA. Over many trials, the response shifts from the delivery of reward (at *t* = 60) to the onset of stimulus (at *t* = 10, dashed line). **(b)** Hypothetical cortical neurons encode time since stimulus onset with a set of temporal tuning curves, as has been suggested [32]. **(c)** Thus, on each trial, the cortical neurons exhibit a cascade of activity. **(d)** We use TD learning theory to constrain a state space model for the activity of cortex and VTA, whose graphical model is shown here (rewards omitted). The latent states are the weights relating cortical activity to an unobserved value function. **(e)** The posterior mean of the latent states of the TD learning state space model. Though not particularly insightful on their own, when combined with cortical activity, the weights determine the posterior distribution of the value function **(f)**. Colors correspond to trials 1, 30, and 150, as in (a). Dotted black line: ground truth. **(g)** We also learn the learning rate, α_l_, under two different models: a constant model and a power-law decay model. **(h)** In contrast to the TD-LDS, fitting a standard LDS to the VTA activity yields accurate predictions, but its latent states are uninformative and do not correspond to weights of a value function.

## Model Criticism and Comparison

Bayesian inference is not the end of the scientific process, but rather an intermediate step in the iterative loop of hypothesizing, fitting, criticizing, and revising a model. Still, posterior inference provides a rigorous and quantifiable method of guiding model criticism and revision. Intuitively, if the model is a good match for the data, then samples from the fit model should “look like” the observed data. *Posterior predictive checks* (PPC’s) [1, 2, 44], which are essentially Bayesian goodness-of-fit tests, formalize this intuition in a statistically rigorous manner. Our presentation here parallels that of Blei [2].

PPC’s compare the observed data to datasets sampled from the posterior predictive distribution (8) of the model. If the sampled data differs from the observed along important dimensions, the model fails the PPC. These “important dimensions” are determined by the practitioner’s choice of a test statistic, *T(x)*: a function that identifies a particular aspect of the data, *x*. For example, in our TD learning simulations, a salient characteristic is the propagation of error signal from the onset of reward to the presentation of the cue. Thus, a simple statistic is amplitude of the error signal in particular trials and time bins. The PPC is defined as the probability that the test statistic of sampled data exceeds that of observed data PPC = Pr(*T*(*x*) > *T*(*x*)| *x*^*^).

The choice of test statistic is left to the practitioner. Clearly, probabilistic modeling under computational constraints necessitates trade-offs and assumptions; no model is perfect. PPC’s are a diagnostic tool for assessing whether the model recapitulates salient features of the data, as determined by the practitioner. In this sense, PPC’s provide a targeted means of criticizing models, shining spotlights on the most important parts. Moreover, there is no limit to the number of PPC’s that may be applied, and the marginal cost of estimating multiple PPC’s is negligible since they can all be estimated using the same sampled data.

Figure 3 illustrates a very simple posterior predictive check for the TD learning model. Panels (a-c) show the observed data (black) and the quantiles of the posterior predictive distribution for the tenth trial, estimated with 1000 samples from the posterior predictive distributions. In this case, the true model uses a power law learning rate, and indeed this is the only model that consistently captures the data. The constant model overestimates the response to the reward (time 60) and the standard LDS incorrectly predicts a response at cue onset. We quantify this with PPC’s for the simplest statistics, *T*_*l,t*_= *x*_*l,t*_.Panels (d-f) show the PPC’s for each trial and time bin. This reveals the delayed responses of the constant model in early trials, and the tendency of the standard LDS to predict a response at cue onset regardless of trial. Under the true model, these PPC’s are uniformly distributed on [0, 1]. Panels (g-f) show that only the power law achieves this.

**Figure 3:**
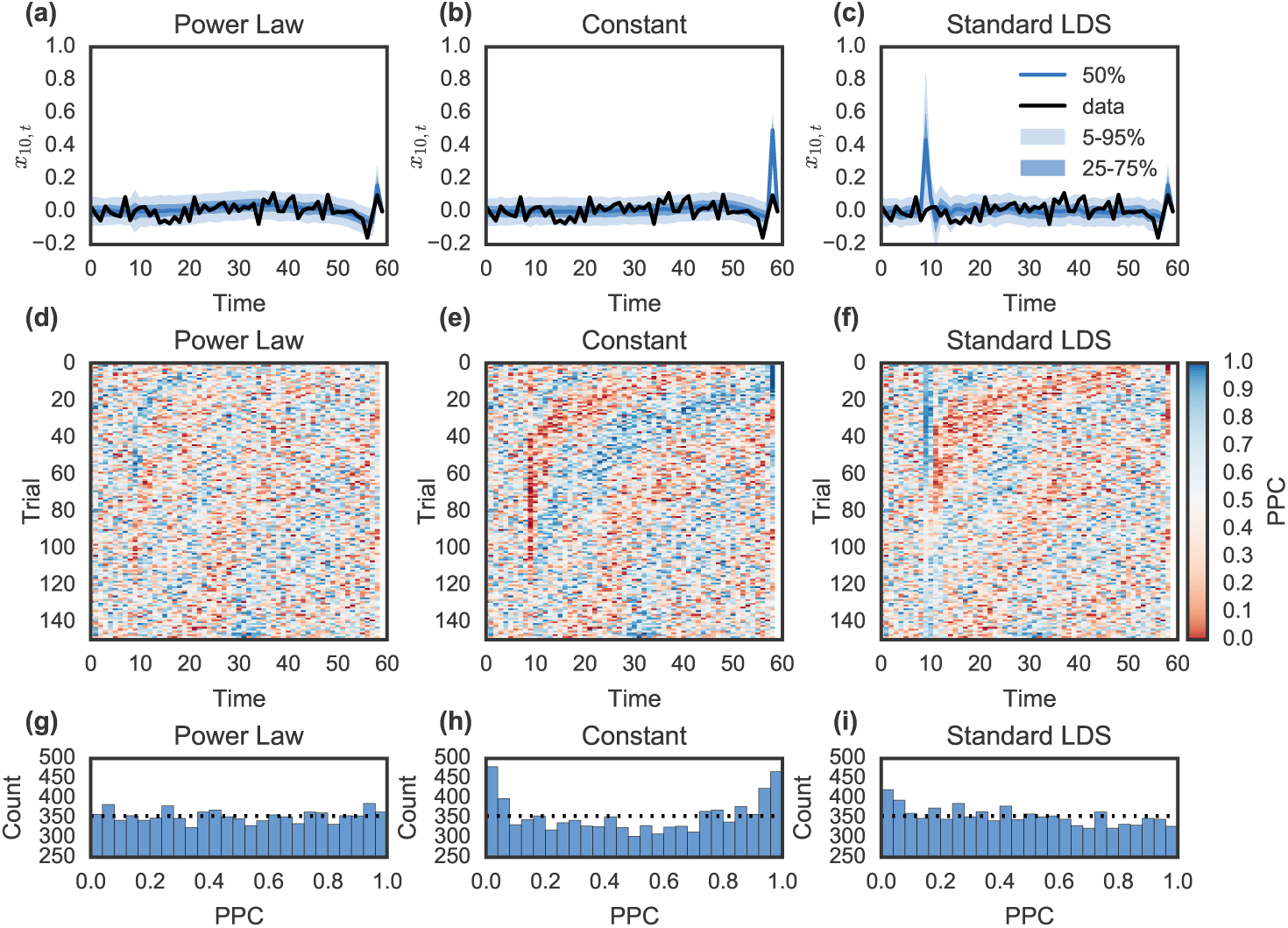
Model criticism using posterior predictive checks (PPC’s). **(a-c)** PPC of the data on trial 10 for three models: the TD-LDS with a power-law learning schedule (i.e. the true model that generated the data); the TD-LDS with a constant learning rate; and a standard LDS. Blue line: posterior predictive median; blue shading: posterior predictive quantiles; black line: observed data. The constant learning rate fails the PPC because it generates a much larger prediction error at time *t* = 59. The standard LDS fails because it always predicts large signals at *t* = 10, regardless of trial. **(d-f)** A summary view of the PPC for all trials and time points. Color denotes the PPC value estimated from 1000 generated trajectories. Blue: model predictions larger than data; red: data larger than model predictions. Values close to zero or one indicate model mismatch. **(g-i)** A histogram of values in (d-f), respectively. The true model should yield uniformly distributed PPC’s(dotted line), as indeed the power law does. The other models generated data that systematically differs from the true data.

While PPC’s check, in absolute terms, how well the model fits the data, in some cases we seek a relative comparison of two models instead. For example, we often cascade models of increasing complexity—factor analysis is a special case of an LDS, which in turn is a special case of a switching LDS—and we need means of justifying this increased capacity. The most straightforward approach is to measure predictive likelihood on held-out data. A better model should assign higher posterior predictive probability, *p*(*x*^*^|*x*), to the held-out data. We see that the predictive probability (8) is an expectation with respect to the posterior. Since this is typically intractable, we estimate the predictive probability with samples from the approximate posterior.

This is by no means the only method of comparing models. In “fully Bayesian” analyses, it is common to compare models on the basis of their marginal likelihood, *p*(*x*) [45, 46]. Recall that this is the denominator in Bayes’ rule (7), and it is generally intractable. Variational methods provide a lower bound on this quantity, and Monte Carlo estimates like annealed importance sampling [47] can yield unbiased estimates of it. In general, however, marginal likelihood estimation is an active area of research [48, 49, 50].

Model criticism suggests not only new theories to test, but also new experiments to run. Specifically, we should choose an experiment that is most likely to reduce the uncertainty of the posterior. Equivalently, we should perform the experiment that yields the maximal information gain in expectation. This intuition is the basis of Bayesian optimal experimental design [45, 51, 52, 53] and is also the guiding principle underlying Bayesian optimization [54]. In our working example, these methods could suggest the combination of stimulus and reward patterns that would be most informative of the underlying learning rate. These methods have been proposed for sampling the voltage on dendritic trees in high-noise settings [55], as well as for designing training regimes for animals [56].

Just as probabilistic programming languages and automated inference algorithms are relieving the burden of Bayesian inference, recent work has attempted to automate model criticism and model comparison. Automatic two-sample tests [57, 58] search for test statistics that best discriminate between the observed data and a model’s samples. In this sense, these approaches are similar to generative adversarial networks [59], which simultaneously train competing generator and discriminator networks. Likewise, automatic model composition methods [60, 61] iteratively construct models, adding increasingly sophisticated structure to capture nuances of the data and comparing on the basis of marginal likelihood. While these advances have still not taken the human “out of the loop,” recent work suggests that these approaches do indeed mimic the process by which humans learn the complex structure of data [62].

## Conclusions

The idea of combining statistical models with computational theories is not new (see [63]), but researchers are only beginning to appreciate the range of possibilities that have opened up with advances in probabilistic modeling. Richly expressive probabilistic programming languages, efficient inference algorithms, and flexible Bayesian nonparametric priors allow complex models to be specified and fit to data much more easily than in the past. Model criticism and comparison techniques can be used to guide the refinement of modeling assumptions, as in Box’s loop. We have shown how this statistical toolbox can be seamlessly integrated with computational theory, using a worked example from reinforcement learning. The key lesson from this modeling exercise is that data-driven and theory-driven approaches to neuroscience need not be mutually exclusive; indeed, the most powerful insights can be gained by using computational theories as constraints on data-driven statistical models.

Conversely, flexible statistical models can enrich computational theories. Historically, computational tractability has biased the kinds of models we fit towards simplicity (conjugacy, convex optimization problems, unimodal posteriors, low-dimensional parametrizations). With faster computers, larger datasets and new algorithms, machine learning has increasingly pushed the envelope towards much more complex models [64, 29, 65], altering the usual tradeoff between neuroscientific realism and computational tractability. We are now in a position to start experimentally testing a vast range of computational theories.

## Acknowledgments

SWL is supported by a Simons Collaboration on the Global Brain postdoctoral fellowship (SCGB-418011). SJG is supported by the National Institutes of Health (CRCNS 1207833). We thank Liam Paninski for his helpful feedback on this work.

## Footnotes

1 Code to run this example and reproduce Figures 2 and 3 is available at https://github.com/slinderman/tdlds

2 We denote the weights by *z* instead of something more traditional, like *w*, since this will highlight the connection to state space models

